# SPARK: A Systems-level Computational Framework for Reconstructing Transcriptomic State Organisation in Lung Adenocarcinoma

**DOI:** 10.64898/2026.06.08.730929

**Authors:** Rishabh Kulkarni, Avik Sengupta, Rahul Kumar

## Abstract

Lung adenocarcinoma (LUAD) exhibits substantial molecular heterogeneity, which complicates tumour stratification and limits the ability of mutation-centric models to capture tumour behaviour and predict patient outcomes. This study investigates whether coordinated transcriptomic programs can provide a systems-level representation of tumour states. Bulk RNA-sequencing data from the TCGA-LUAD cohort were analysed to reconstruct pathway-level transcriptomic organisation using a stability-optimised network framework (SPARK). This analysis identified eight transcriptomic modules representing coordinated biological processes active across tumours.

Module activity scores were subsequently used to derive a composite Transcriptomic Risk Score through elastic-net Cox proportional hazards modelling. The resulting risk score showed a significant association with overall survival in the discovery cohort and improved prognostic discrimination beyond clinical variables. An independent evaluation in the CPTAC-LUAD cohort confirmed the prognostic signal and preserved risk stratification across patient groups. Unsupervised clustering of module activity further revealed three transcriptomic patient groups characterised by distinct biological programs, genomic alteration patterns, and survival outcomes. Single-cell analysis also demonstrated that the identified transcriptomic modules reflect coordinated organisation of the tumour–immune–stromal ecosystem across cellular compartments.

Together, these findings suggest that LUAD heterogeneity can be organised into coordinated transcriptomic programs with measurable clinical relevance, providing a systems-level framework for representing tumour molecular states.

## Introduction

Lung adenocarcinoma (LUAD) is characterised by substantial molecular diversity, with recurrent alterations in key driver genes such as *EGFR, KRAS*, and *TP53* contributing to variability in tumour behaviour and clinical outcomes [1]. Multiple transcriptomic profiling studies have also revealed variation in pathway activity across tumours, including proliferation, metabolism, immune signalling, and microenvironmental interactions. However, these molecular signals often coexist within the same tumour, making it difficult to integrate them into a coherent representation of tumour biology. Therefore, translating high-dimensional transcriptomic variation into clinically meaningful insights remains a central challenge in LUAD and in cancer biology more broadly [2].

Representing tumour behaviour through individual genes or multi-gene signatures presents several challenges. Gene expression is subject to biological variability and technical noise, making such measurements difficult to interpret [3]. Multi-gene signatures attempt to address this limitation by combining gene sets to develop predictive or prognostic models. However, these approaches represent limited aspects of tumour biology and neglect broader molecular context, resulting in weak reproducibility across independent cohorts [4]. These limitations highlight the need for broader representations of transcriptomic activity in tumours.

Biological pathways provide one such representation for modelling tumour states, as they group genes involved in shared cellular processes [5, 6] However, pathways are often highly interconnected and overlapping, with many genes participating in multiple biological processes. Therefore, tumour states may be more effectively described through higher-order transcriptomic modules that summarise coordinated activity across multiple pathways [7]. These modules capture dominant patterns of biological activity within tumours and provide compact, low-dimensional representations while preserving biologically meaningful information.

Here, we developed SPARK (a graph-based stability-optimised program architecture reconstruction framework) to identify reproducible modules of coordinated pathway activity. Using treatment-naïve patients from the TCGA-LUAD cohort, we define a low-dimensional representation of tumours based on module activity, capturing tumour variation along a continuous transcriptomic axis. Regularised survival modelling was used to derive a composite risk score associated with patient outcomes, and validated generalisability was evaluated in an independent cohort from the CPTAC-LUAD. An overview of the study design and analytical workflow is shown in **Figure 1A**.

**Figure 1.**
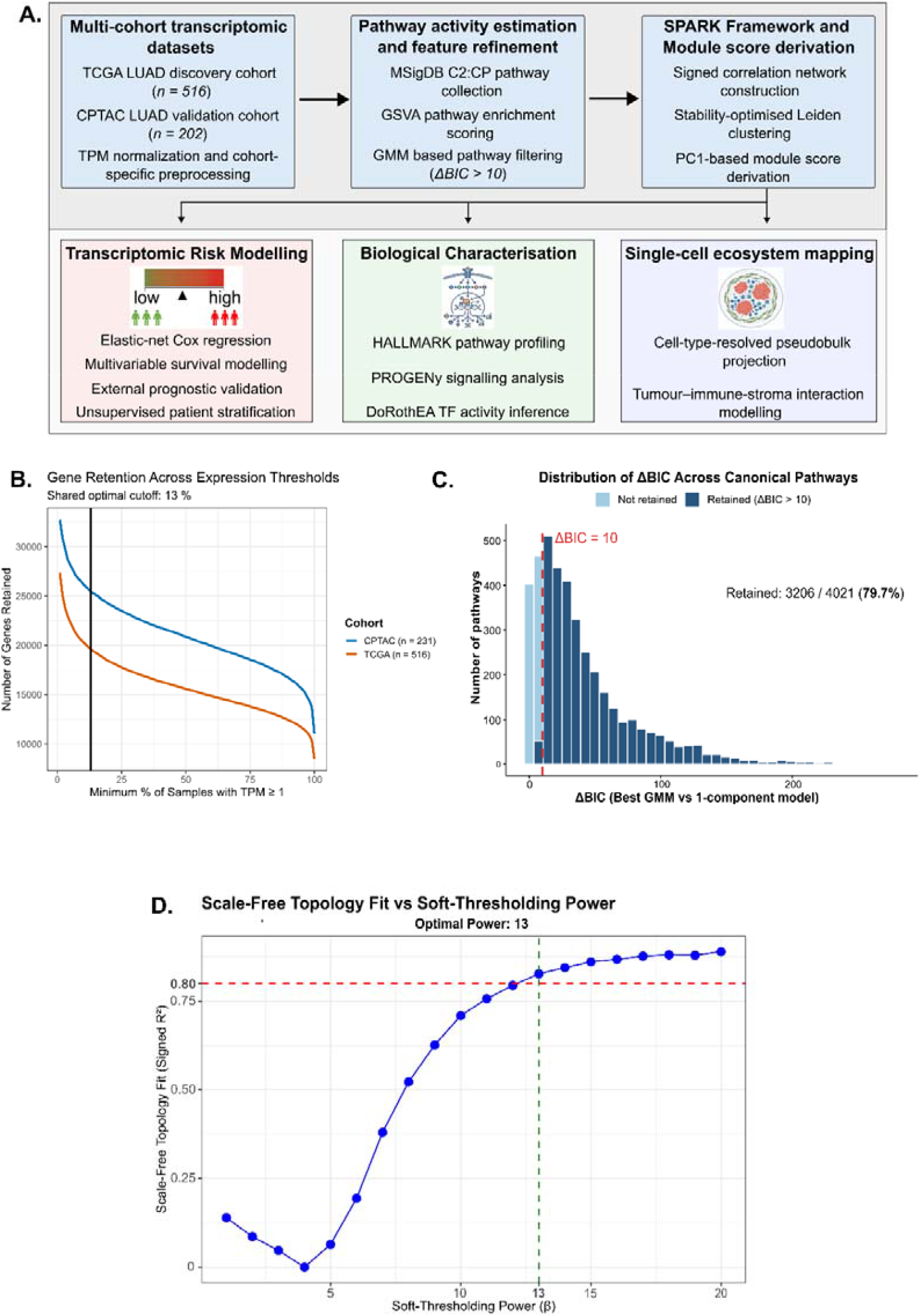
Cohort overview, transcriptomic pre-processing and pathway feature refinement. **(A)** Overview of the analytical framework integrating bulk transcriptomic modelling, SPARK-based module reconstruction, transcriptomic risk modelling, biological characterisation, and single-cell ecosystem analysis across LUAD cohorts. **(B)** Gene retention curves across expression thresholds in the TCGA LUAD and CPTAC LUAD cohorts. The elbow heuristic identified an optimal filtering threshold of 13% of samples with TPM ≥ 1. **(C)** Distribution of ΔBIC values comparing Gaussian mixture models with single-component models across canonical pathways. Pathways with ΔBIC > 10 were retained for downstream module reconstruction, yielding 3206 retained pathways (79.7%). **(D)** Scale-free topology fits across candidate soft-thresholding powers for pathway–pathway correlation network construction. The selected soft-thresholding power achieved a scale-free topology index > 0.8.

## Methods

### Cohorts and Transcriptomic Processing

Transcriptomic and clinical data for Lung Adenocarcinoma (LUAD) were obtained from the cBioPortal platform, using the Genomic Data Commons (GDC) release of The Cancer Genome Atlas (TCGA) LUAD cohort and the Clinical Proteomic Tumour Analysis Consortium (CPTAC) LUAD cohort [8, 9]. Only primary tumours with matched RNA-seq and clinical annotations were retained. Patients without overall survival (OS) information were excluded.

Gene expression data were obtained in Transcripts per million (TPM) format. Low-expression genes were filtered using a minimum threshold of TPM ≥ 1 in a cohort-specific proportion of samples, determined using a data-driven elbow-based heuristic. Expression values were log-transformed (log_2_(TPM + 1)) for downstream analysis. All preprocessing steps were applied consistently across cohorts to ensure comparability and preserve downstream model transferability.

### Pathway Activity Estimation

Pathway activity scores were estimated using the Molecular Signatures Database (MSigDB) C2: CP (curated canonical pathways) gene set collection (version 2025.1, Homo sapiens) [5]. Patient-wise pathway enrichment scores were estimated using Gene Set Variation Analysis (GSVA) implemented in the GSVA package in R [10]. Expression matrices were normalised using log2(TPM + 1), and a Gaussian kernel cumulative distribution function was applied, appropriate for continuous RNA-seq data. The resulting output was a pathway activity matrix of dimension *P × N*, where *P* denotes the number of C2: CP pathways and *N* the number of patient samples per cohort.

### Stability-optimised Program Architecture Reconstruction Framework (SPARK)

A higher-order transcriptomic organisation (modules) was formalised from pathway activity profiles using the Stability-optimised Program Architecture Reconstruction Framework (SPARK), applied to the TCGA LUAD cohort. SPARK integrates multimodality filtering, network construction, community detection, and stability-based parameter selection within a reproducible workflow.

Many clustering approaches used to identify structure in biological networks produce a single partition of the data and have limited ability to evaluate multiple community resolutions. However, complex biological networks usually exhibit community structure across multiple topological scales [11]. SPARK was therefore designed to enable a systematic exploration of the pathway network organisation and identify a stable community partition.

### Multimodality Filtering

Pathways were filtered by fitting Gaussian Mixture Models (GMMs) to GSVA score distributions using the mclust package in R [12]. Models with varying numbers of components were evaluated, and selection was performed using the Bayesian Information Criterion (BIC).

For each pathway, ΔBIC was defined as:

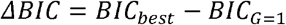

where *BIC*_*best*_ corresponds to the optimal mixture model, and *BICG=1* corresponds to the single-component (unimodal) model. Larger ΔBIC values indicate stronger evidence in favour of multimodality. Following established guidelines for BIC differences, pathways with Δ**BIC > 10** -based module discovery [13].

### Correlation network construction

A signed pathway–pathway correlation network was constructed using the retained GSVA pathway scores. Pearson correlation coefficients were calculated between all pairs of pathways across patient samples. To approximate a scale-free topology in the network, a soft-thresholding power *β* was selected using the WGCNA framework [14]d between 1 and 20 (step size = 1), under the signed network framework. For each *β*, the scale-free topology index (R2) was computed. The optimal power *β*_*opt*_ was chosen as the smallest power that achieved a scale-free topology fit index (**R**^**2**^ **> 0.8**).

The adjacency matrix *Aij*, representing pairwise pathway correlation, was defined using the signed power function:

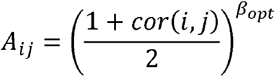

where *cor(i,j)* denotes the Pearson correlation between pathways *i* and *j*, and *βopt* is the selected soft-thresholding power. A weighted undirected graph was constructed from the adjacency matrix using the igraph package in R [15], where nodes represent pathways and edge weights correspond to the adjacency values.

### Community Detection

Community detection was performed using the Leiden algorithm [16], which was implemented in the igraph package in R. The algorithm optimally partitions a weighted graph into communities by maximising the modularity.

For a weighted undirected graph with adjacency matrix *Aij*, modularity is defined as:

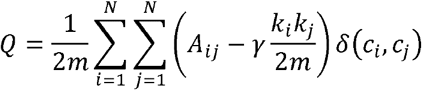

where,

- *Aij* denotes the edge weight between pathways *i* and *j*,
- 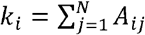 is the weighted degree of node *i*,
- 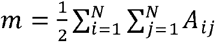 is the total edge weight in the graph,
- γ is the resolution parameter,
- *ci* is the community assignment of node *i*,
- δ (ci, cj) = 1 if nodes *i* and *j* are in the same community and 0 otherwise.

The algorithm iteratively optimises Q through local node movements and community refinement, guaranteeing well-connected communities.

The weighted pathway graph was partitioned across a resolution parameter grid ranging from 0.5 to 2.0 (step size 0.1). Full convergence was enforced (*niterations = -1*). For each resolution value, pathways were assigned to modules, and modules containing fewer than two pathways were excluded from downstream module scoring.

### Stability-based resolution selection

The optimal resolution parameter was determined by evaluating clustering stability and within-module correlation structure across the resolution grid:

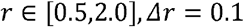

For each resolution value *r*, bootstrap stability was assessed using 500 iterations (n_boot_). In each iteration, two independent resampling procedures were performed. First, Pathway resampling: 80% of pathways were sampled without replacement, and Leiden clustering was repeated on the induced subgraph. Second, Sample resampling: 80% of patient samples were resampled without replacement; a new adjacency matrix was recomputed with the same soft-thresholding power *β*; and Leiden clustering was repeated. For each bootstrap iteration, concordance between the original and bootstrap clustering was quantified using the Adjusted Rand Index (ARI) [17], computed with the mclust package. Two ARI values were recorded per iteration: pathway-level and sample-level ARI. Stability (*GeoStability*) was defined as the geometric mean of the mean pathway-level ARI and mean sample-level ARI for every resolution *r*:

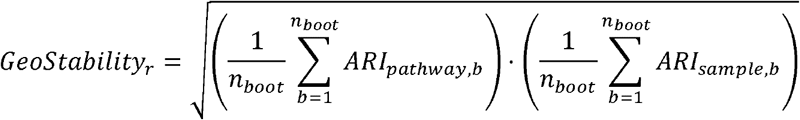

Module-level correlation structure was quantified using the mean absolute pairwise correlation (MAPC). For a module *m* containing *km* pathways, MAPC was computed as:

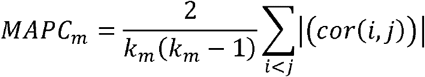

where correlations were computed from the GSVA score matrix, for each resolution, a weighted average MAPC was calculated using module sizes as weights:

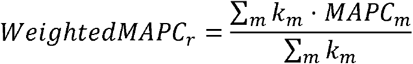

To integrate stability and correlation structure, metrics were optimised across resolutions using min–max scaling. For a metric *Xr* evaluated at resolution *r*, normalisation was defined as:

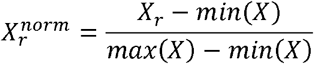

Entropy-based weights (*w*_*j*_) were computed from the normalised metric matrix:

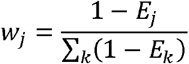

where *Ej* denotes the normalised entropy of metric *j*.

Finally, the composite score was defined as:

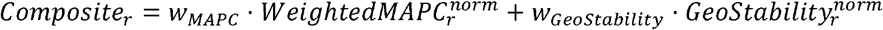

The optimal resolution parameter *ropt* was selected as the value *r* that maximises the composite score. In summary, SPARK integrates GMM (mclust), network construction (WGCNA, igraph), Leiden community detection (igraph), and entropy-weighted stability optimisation within a reproducible computational workflow.

### Module Score Construction

Pathway modules identified by SPARK were summarised into quantitative features using principal component analysis (PCA). For each module *m*, the corresponding pathway activity matrix was centred and scaled, and the first principal component (PC1) loading vector *wm* was extracted to represent the dominant axis of coordinated pathway variation.

For each patient *i*, the module score was defined as the projection of pathway activity onto this axis:

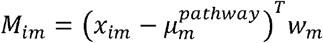

where *x*_*im*_ denotes the pathway activity vector and μ_*m*_ the pathway-wise mean estimated from the discovery cohort.

Module scores were standardised within the discovery cohort, and the derived loading vectors and scaling parameters were retained as fixed bases. External cohort scores were computed by projection onto these fixed bases without re-estimation, ensuring consistency across datasets.

### Unsupervised patient stratification

Unsupervised clustering was performed on the standardised module score matrix using the ConsensusClusterPlus package in R [18]. Hierarchical clustering was evaluated across a range of cluster numbers (K = 2–20) using multiple distance metrics (Euclidean, Pearson, and Spearman) and linkage methods (Ward.D2, average, and complete). For each configuration, clustering was performed over 1000 resampling iterations, with 80% of patients subsampled without replacement per iteration while retaining all module features. Clustering stability for each candidate K was evaluated using multiple complementary metrics, including the proportion of ambiguous clustering (PAC), mean silhouette width, changes in the consensus cumulative distribution function (ΔCDF), and mean cluster consensus. These metrics were min–max normalised and integrated using an entropy-weighted composite score to identify the most stable K within each distance-linkage configuration. Concordance between clustering solutions across configurations was additionally assessed using the Adjusted Rand Index (ARI). Among the highest-ranking stable clustering solutions, the final configuration was selected based on downstream biological relevance, including survival separation across patient groups. The final solution corresponded to Euclidean distance with Ward.D2 linkage and K = 3 clusters.

### Transcriptomic Risk Score Derivation

A transcriptomic risk score was derived using an elastic net penalised Cox proportional hazards model with module scores as predictors. Overall survival was defined as the time from diagnosis to death or last follow-up, with right censoring. Model fitting was performed using the survival and glmnet R packages [19, 20].

Hyperparameters were selected using nested cross-validation, with a 5-fold outer loop and 10-fold inner loop. The mixing parameter (α) was optimised over a predefined grid, and the regularisation parameter (λ) was selected based on cross-validated deviance. Model performance was evaluated using the concordance index (C-index), and optimal parameters were chosen based on mean outer-fold performance [21]. The final model was refit on the full discovery cohort using the selected parameters. Feature robustness was further evaluated using bootstrap stability selection (200 iterations), in which the model was refitted across bootstrap resamples to quantify module selection frequency [22]. Variable importance was additionally assessed through permutation testing by measuring reductions in model C-index following module-wise score randomisation.

The risk score for patient *i* was defined as:

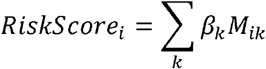

where *β*_*k*_ are model coefficients, and *M*_*ik*_ denotes the score of module *k* for patient *i*.

### Biological characterisation of transcriptomic modules

Functional annotation of module scores was carried out using pathway and transcription factor activity inference frameworks applied to log2(TPM+1) expression data. Hallmark pathway activity (MSigDB v2025.1. Hs) was estimated using the GSVA package, generating a sample × pathway activity matrix. Transcription factor activity was inferred using DoRothEA human regulons (confidence levels A–C) implemented in the decoupleR framework in R, which uses the VIPER algorithm [23]. Signalling pathway activity was additionally quantified using the progeny package in R [24]. Module scores were subjected to an independent Pearson correlation analysis with all the calculated activities. P-values were adjusted using the Benjamini–Hochberg false discovery rate procedure (p-adj < 0.05).

### Single-cell analysis

Single-cell RNA-seq data from a primary LUAD cohort (GSE131907; n = 15 patients, 57,155 cells) were used to evaluate module activity at the single-cell level [25]. Cells were annotated into tumour (epithelial), immune, and stromal compartments, and cell-type–resolved pseudobulk profiles were generated for each compartment within each patient. Pathway activity scores were computed using the same GSVA framework as in bulk data and projected onto SPARK-derived modules using fixed loading vectors. Module activity was summarised as compartment-level z-scores, and tumour–immune interaction states were defined based on the sign of tumour and immune scores. Stromal activity was analysed in relation to tumour–immune balance to assess context-dependent coordination across compartments.

## Results

### Cohort characteristics and transcriptomic preprocessing

The TCGA LUAD cohort was used as the discovery dataset (n = 516 patients, with complete transcriptomic, clinical, and survival annotations), and the CPTAC LUAD cohort as an external validation dataset (n = 231 patients, 202 with complete transcriptomic, clinical, and survival annotations). Detailed cohort characteristics for both cohorts are provided in **Table 1**.

**Table 1.**
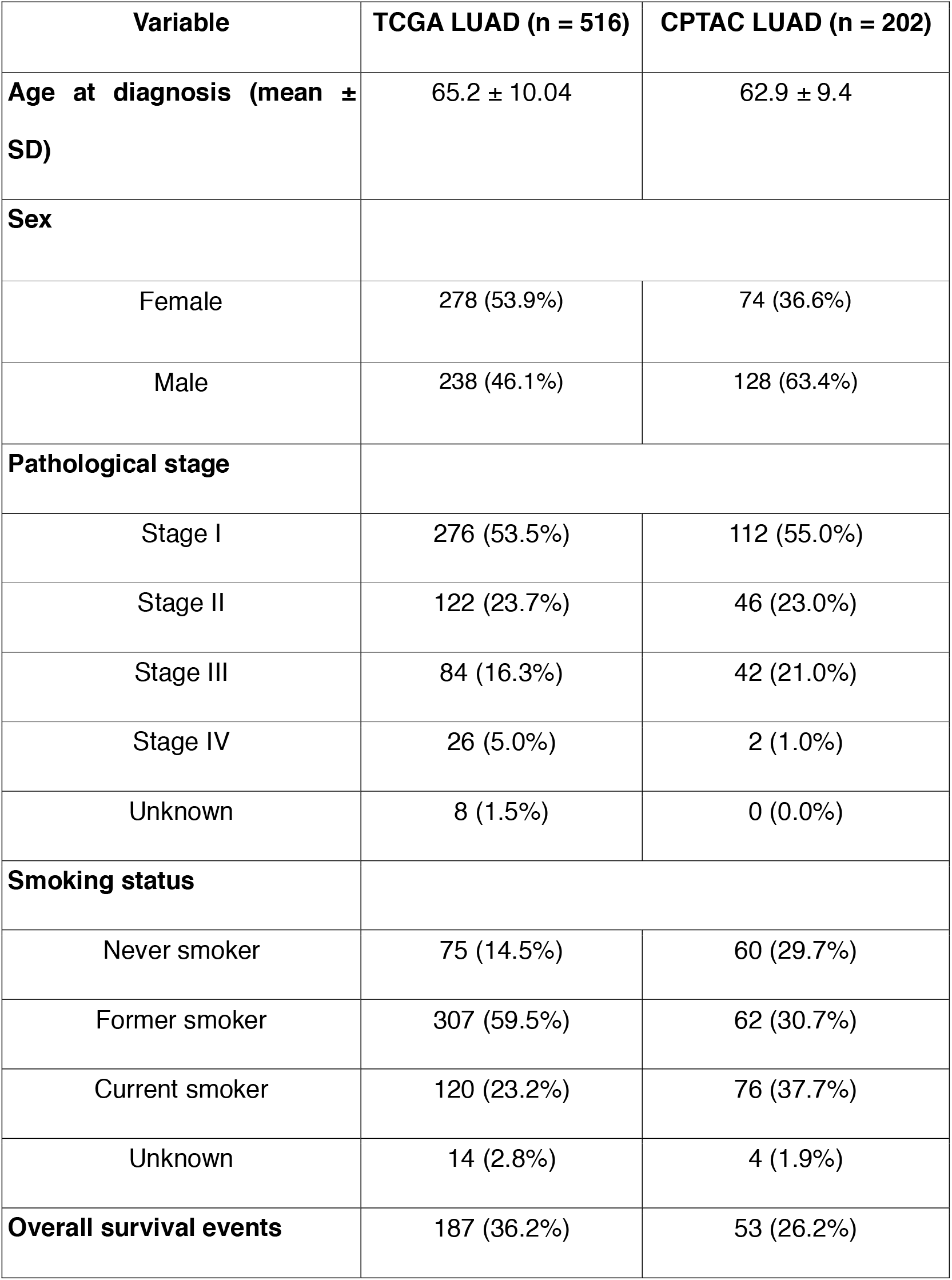
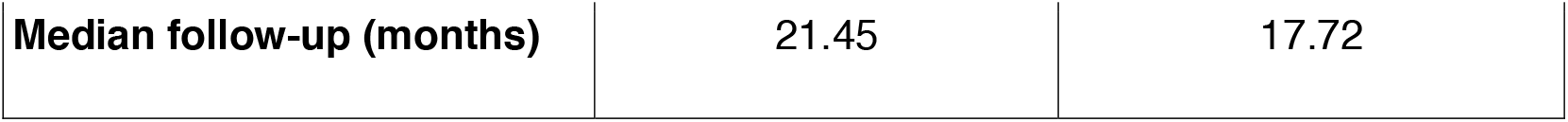
Baseline clinical characteristics of the TCGA LUAD discovery cohort and CPTAC LUAD validation cohort.

| Variable | TCGA LUAD (n = 516) | CPTAC LUAD (n = 202) |
| --- | --- | --- |
| Age at diagnosis (mean ± SD) | 65.2 ± 10.04 | 62.9 ± 9.4 |
| Sex |  |  |
| Female | 278 (53.9%) | 74 (36.6%) |
| Male | 238 (46.1%) | 128 (63.4%) |
| Pathological stage |  |  |
| Stage I | 276 (53.5%) | 112 (55.0%) |
| Stage II | 122 (23.7%) | 46 (23.0%) |
| Stage III | 84 (16.3%) | 42 (21.0%) |
| Stage IV | 26 (5.0%) | 2 (1.0%) |
| Unknown | 8 (1.5%) | 0 (0.0%) |
| Smoking status |  |  |
| Never smoker | 75 (14.5%) | 60 (29.7%) |
| Former smoker | 307 (59.5%) | 62 (30.7%) |
| Current smoker | 120 (23.2%) | 76 (37.7%) |
| Unknown | 14 (2.8%) | 4 (1.9%) |
| Overall survival events | 187 (36.2%) | 53 (26.2%) |
| **Median follow-up (months)** | 21.45 | 17.72 |

Transcriptomic data was processed to ensure consistency across cohorts. Following gene symbol harmonisation, over 40,000 genes were retained, and low-expression genes were filtered using an elbow-curve-based minimum expression threshold (TPM ≥ 1 in at least 13% of samples), yielding 19,595 genes in the TCGA cohort and 25,452 in the CPTAC cohort for downstream analysis (**Figure 1B**).

Pathway activity scores were estimated using gene set variation analysis across 4021 canonical pathways. To identify pathways exhibiting heterogeneous activity across patients, Gaussian mixture models were applied, and pathways with ΔBIC > 10 were retained. This resulted in 3206 multimodal pathways, which were carried forward for network-based module reconstruction (**Figure 1C**).

### Stable pathway modules define transcriptomic architecture

The Stability-optimised Program Architecture Reconstruction Framework (SPARK) was applied to derive a stable organisation of the 3206 multimodal pathways retained following ΔBIC-based filtering.

A signed pathway–pathway correlation network was constructed using pairwise Pearson correlation coefficients computed across pathway activities in the discovery cohort. To approximate scale-free topology, candidate soft-thresholding powers (*β*) were evaluated, and the smallest power (*β*_opt_ = 13) leading to a scale-free topology index greater than 0.8 was selected (**Figure 1D**). The resulting adjacency matrix comprised 3,206 nodes and 5,137,615 weighted edges and was used for downstream community detection.

The network was partitioned using the Leiden algorithm across a resolution grid. The number of detected modules increased progressively with resolution, ranging from a few at low resolution to higher fragmentation at higher resolutions. To identify an optimal partition, clustering stability and within-module structural coherence were evaluated across the resolution grid and integrated using a composite score (**Figure 2A**).

**Figure 2.**
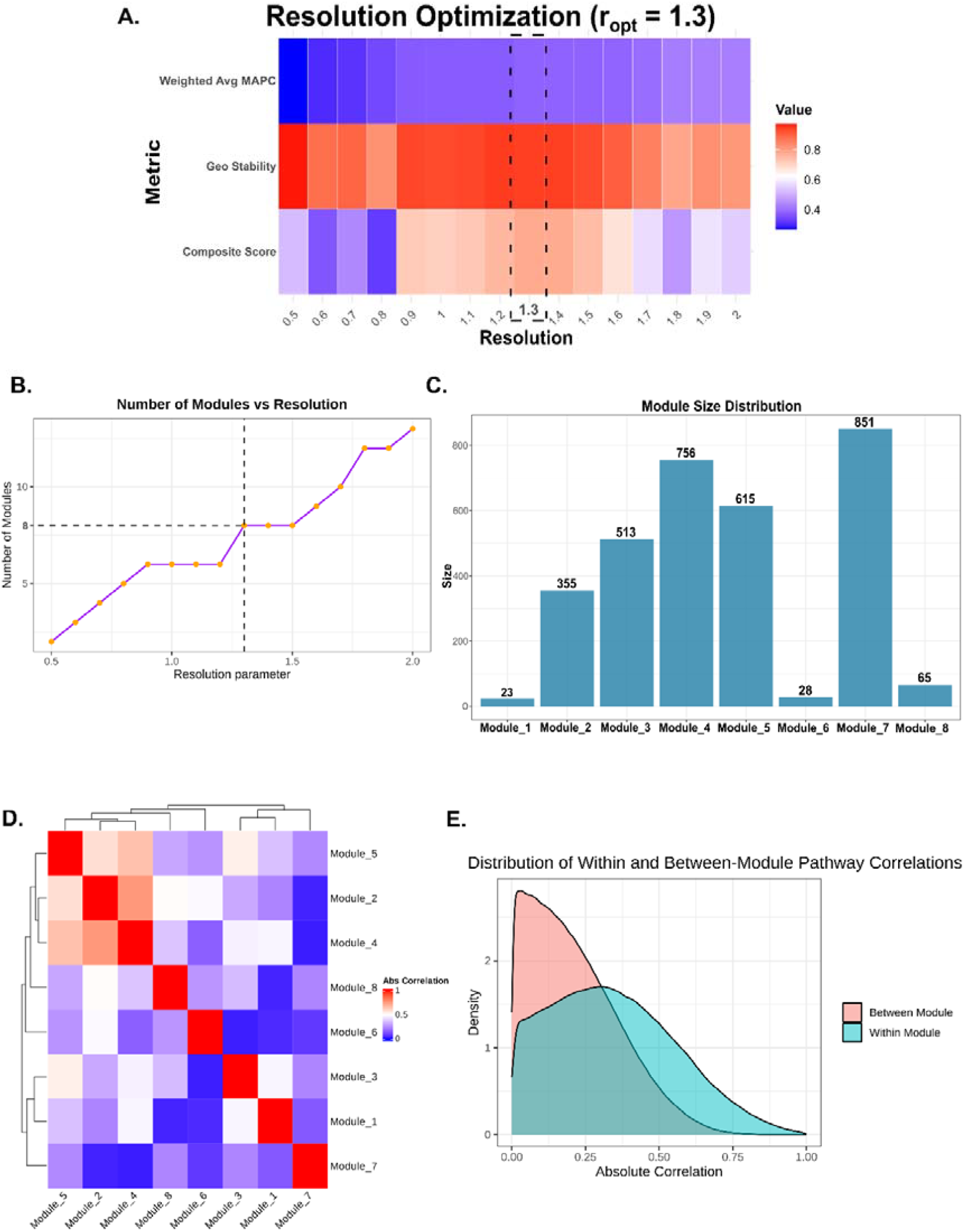
Stable pathway module reconstruction using the SPARK framework. **(A)** Resolution optimisation across Leiden clustering parameters, integrating clustering stability and module coherence. The optimal resolution was identified at r_opt_ = 1.3. **(B)** Number of detected modules across the Leiden resolution parameter grid. **(C)** Distribution of pathway counts across the eight SPARK-derived transcriptomic modules. **(D)** Pairwise correlation heatmap of module activity scores across patients. **(E)** Distribution of within-module and between-module pathway correlations, demonstrating increased coherence among pathways assigned to the same module.

At the optimal resolution (r_opt_ = 1.3), the pathway network was partitioned into eight modules (**Figure 2B**). Module sizes ranged from 23 to 851 pathways, with a median of approximately 464, indicating balanced partitioning without excessive fragmentation or dominance by a single module (**Figure 2C**). Complete resolution-wise optimisation metrics, including GeoStability, weighted MAPC, entropy weights, and composite scores across all tested resolutions, are provided in **Supplementary Table 1**.

### Modules define a low-dimensional tumour state space

Module partitions were transformed into quantitative features via principal component-based projection, yielding a score matrix. These scores define a low-dimensional transcriptomic space, where each tumour is represented as a point defined by module activity.

Pairwise correlation analysis of module activity scores revealed a structured organisation of tumour biology. Correlation coefficients ranged from -0.76 (ρ_min_) to 0.66 (ρ_max_), with a near-zero mean and moderate absolute correlation. No module pair exhibited near-collinearity, indicating that modules capture largely independent axes of variation (**Figure 2D**).

At the pathway level, within-module correlations were consistently higher than between-module correlations, demonstrating that pathways grouped within each module exhibit coordinated behaviour relative to the broader network. This separation between intra- and inter-module correlation distributions points to the structural coherence of the module definitions **(Figure 2E**). Other quantitative characteristics of the modules, including module size, PC1 variance explained and MAPC, are summarised in **Table 2**.

**Table 2.** Quantitative characteristics of the SPARK-derived pathway modules in the TCGA LUAD discovery cohort.

| **Module** | **Number of pathways** | **PC1 variance (%)** | **MAPC** |
| --- | --- | --- | --- |
| 1 | 23 | 92.68 | 0.92 |
| 2 | 355 | 63.9 | 0.62 |
| 3 | 513 | 50.91 | 0.48 |
| 4 | 756 | 25.99 | 0.25 |
| 5 | 615 | 35.93 | 0.33 |
| 6 | 28 | 70.39 | 0.67 |
| 7 | 851 | 35.19 | 0.33 |
| 8 | 65 | 80.86 | 0.79 |

Therefore, this low-dimensional structure provided a compact representation of tumour heterogeneity and served as the basis for downstream prognostic, patient stratification, and single-cell analyses.

### A transcriptomic risk axis emerges from the module-defined state space

Risk scores were derived using an Elastic-Net Cox proportional hazards model with module scores from the TCGA LUAD discovery cohort as predictors. Hyperparameter optimisation was performed within a cross-validated (inner ten-fold and outer five-fold) framework, and model performance was quantified using the C-index metric. The model achieved a mean outer five-fold C-index of 0.613 ± 0.044 (**Figure 3A**). When refitted on the full cohort, a sparse solution with seven non-zero modules was observed (**Figure 3B**), with bootstrap stability analysis showing consistent feature selection across resampling iterations and permutation-based importance analysis confirming that a subset of modules contributed disproportionately to model performance (**Supplementary Figure 1A and 1B**). Further characterisation of the prognostic contribution was performed through module-wise multivariable Cox analyses, which identified a subset of modules with significant associations with overall survival (**Figure 3C**).

**Figure 3.**
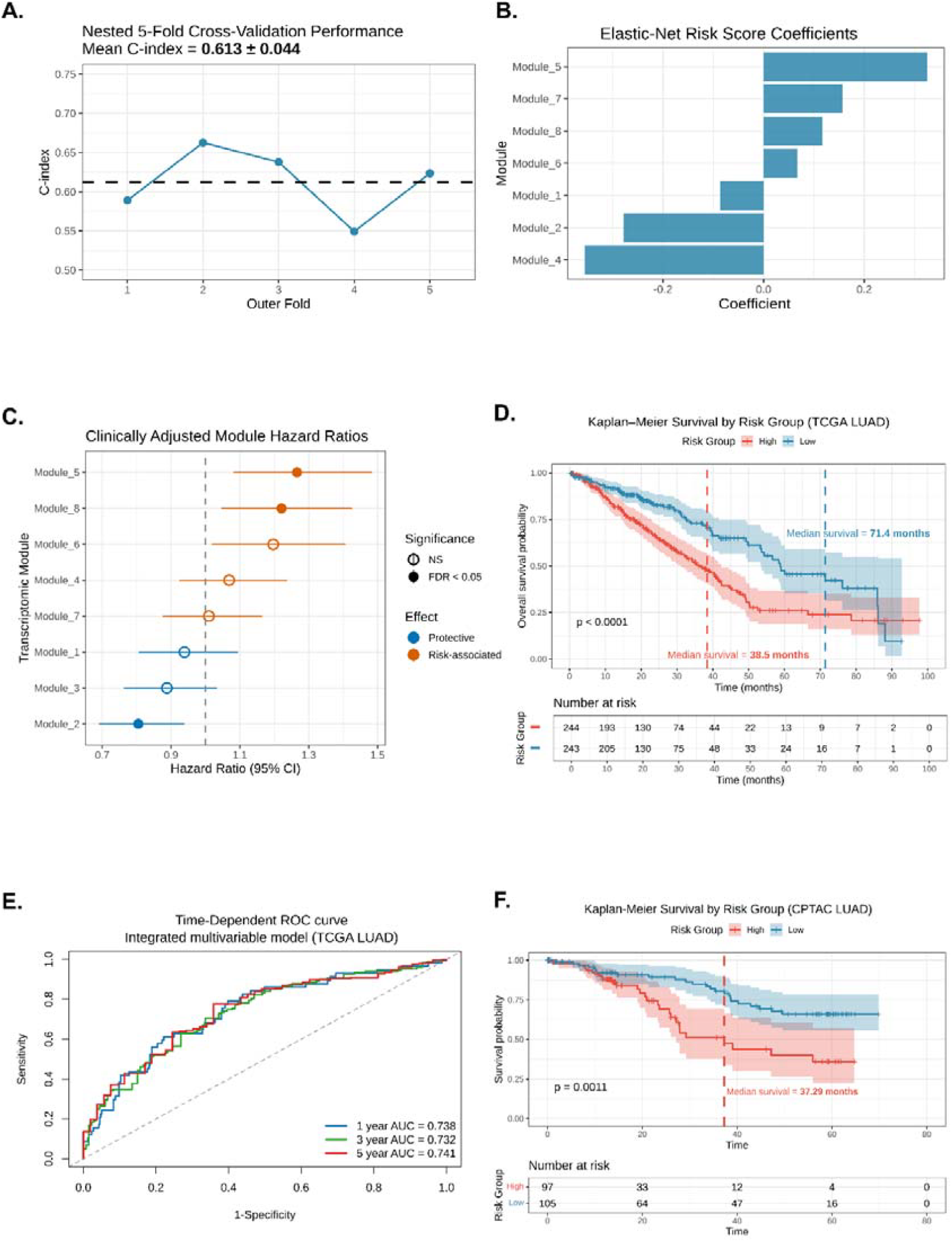
Derivation and validation of the transcriptomic risk axis. **(A)** Nested five-fold cross-validation performance of the Elastic-Net Cox proportional hazards model across outer folds. Mean outer-fold C-index = 0.613 ± 0.044. **(B)** Elastic-Net coefficients for SPARK-derived modules following model refitting on the full TCGA LUAD cohort. **(C)** Multivariable Cox proportional hazards analysis integrating transcriptomic risk score and clinical covariates. **(D)** Kaplan–Meier survival analysis comparing high-risk and low-risk groups in the TCGA LUAD cohort (log-rank p = 1.24 × 10^−^□). **(E)** Time-dependent receiver operating characteristic analysis comparing clinical, transcriptomic, and integrated prognostic models in the TCGA LUAD cohort. **(F)** Kaplan–Meier survival analysis for external validation of the transcriptomic risk score in the CPTAC LUAD cohort (log-rank p = 0.0011).

The distribution of risk scores was approximately unimodal, ranging from -1.01 to 1.27, with a median of 0.011, reflecting patient variability along a continuous risk axis. The cohort median was used as a predefined threshold to stratify patients into high- and low-risk groups. Kaplan–Meier survival analysis demonstrated a highly significant separation between groups (log-rank p = 1.24e-05), with median overall survival of 38.5 months in the high-risk group compared to 71.4 months in the low-risk group (**Figure 3D**).

This separation was also preserved within early-stage disease, with significant survival differences observed among patients with stage I/II tumours, indicating that the transcriptomic axis captures heterogeneity not resolved by pathological staging (**Supplementary Figure 1C**). High-risk tumours showed greater activity of proliferative, growth-metabolic coupling, and genome instability programmes, whereas low-risk tumours were associated with immune-related and tumour suppressor programmes.

Following multivariable Cox regression incorporating clinical covariates, the risk score remained independently associated with survival (HR = 2.66, 95% CI 1.78– 3.98, p = 1.91 × 10^-6^, **Supplementary Table 2**), indicating that it captures prognostic information beyond standard clinicopathologic parameters. The clinical baseline model achieved a mean C-index of 0.688, while the transcriptomic risk score model achieved a mean C-index of 0.635. Integration of the risk score with clinical covariates increased the mean C-index to 0.711, showing incremental prognostic value beyond standard clinicopathologic factors. Additionally, time-dependent receiver operating characteristic (ROC) analysis was performed to evaluate the discrimination power of the final multivariable model across clinically relevant time periods (**Figure 3E**). It achieved an AUC of 0.738 at 1 year, 0.732 at 3 years, and 0.741 at 5 years. This indicates stable, strong predictive performance across early and late follow-up in the cohort.

External validation in the CPTAC LUAD cohort demonstrated consistent generalisation. Risk scores were computed using fixed model parameters without retraining, and significant survival separation between high- and low-risk groups was observed (log-rank p = 0.0011). Discrimination performance was preserved, and integration with clinical covariates improved predictive performance over the clinical baseline (**Figure 3F**). Pathway-to-module assignments, PC1 loading vectors, and module scaling parameters used for projection are provided in **Supplementary Tables 3 and 4**.

### Transcriptomic Modules capture distinct biological programs in LUAD

Functional annotation showed that all SPARK-derived modules captured distinct biological programs associated with LUAD. Modules 4 and 6 were enriched for MYC/E2F-driven proliferative programs, while Modules 1 and 3 captured humoral immune activation and cytokine-mediated inflammatory signalling, respectively. Module 7 was associated with developmental and remodelling processes, including TGF-*β* signalling and epithelial plasticity (**Figure 4A**).

**Figure 4.**
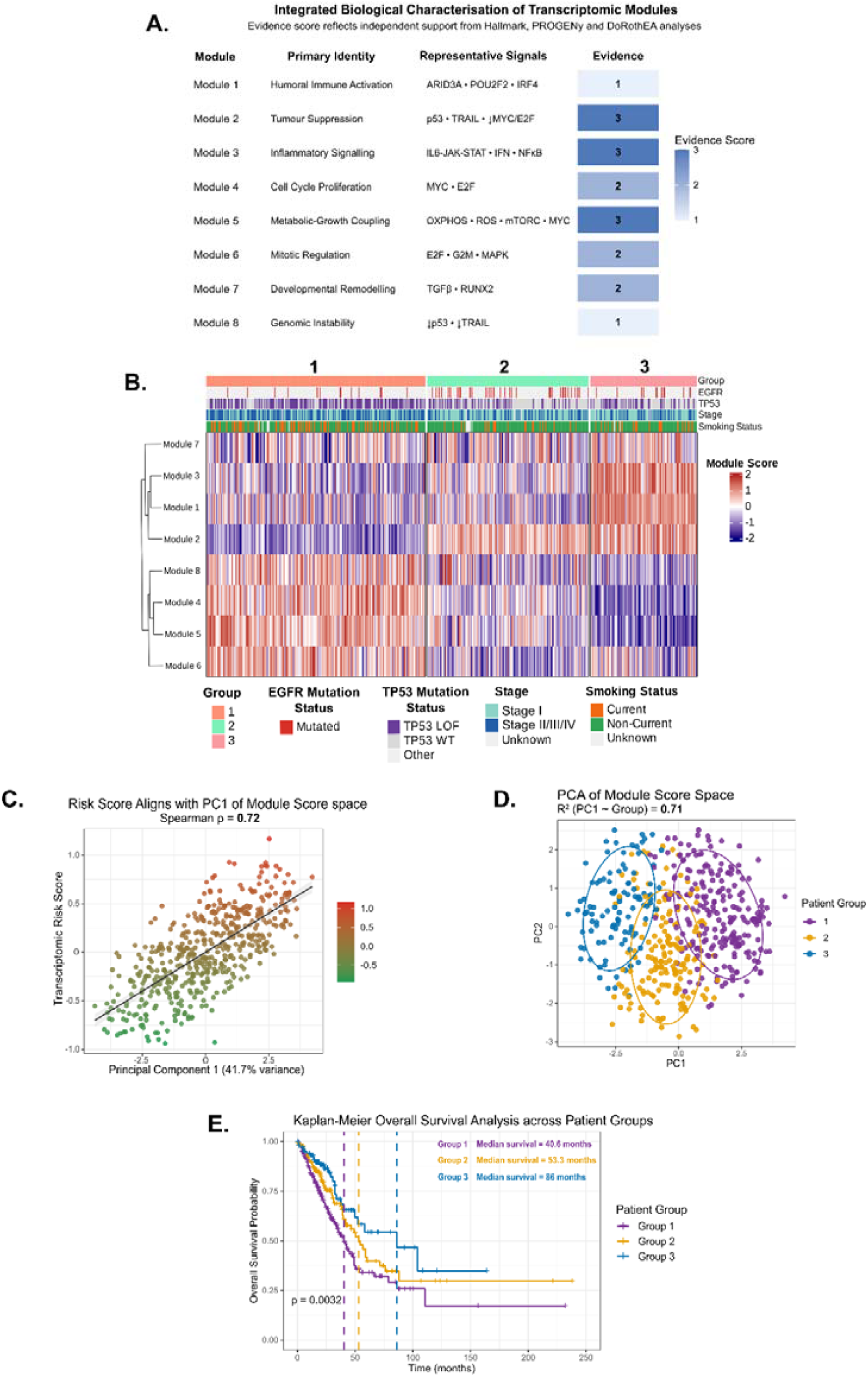
Transcriptomic state space organisation and biologically distinct patient groups. **(A)** Integrated biological characterisation of SPARK-derived transcriptomic modules using Hallmark, PROGENy, and DoRothEA evidence. **(B)** Heatmap of module activity scores across patients grouped by transcriptomic state, annotated with EGFR mutation status, TP53 mutation status, pathological stage, and smoking status. (**C)** Correlation between transcriptomic risk score and the dominant principal component of the module-defined state space (Spearman’s ρ = 0.72). **(D)** Projection of patients into principal component space coloured by transcriptomic state. Patient groups formed distinguishable but partially overlapping regions within the module-defined state space. **(E)** Kaplan–Meier survival analysis across transcriptomic patient groups (log-rank p = 0.0032). Median overall survival was 40.6 months in Group 1, 53.3 months in Group 2, and 86 months in Group 3.

Module-wise multivariable Cox analyses identified Modules 2, 5, and 8 as having the strongest independent associations with overall survival (**Figure 3C**). Functional enrichment revealed that Module 2 represented a tumour suppressor-associated state, whereas Module 5 captured oxidative phosphorylation and mTORC1-associated metabolic signalling. Module 8 was enriched for chromatin remodelling and DNA damage response pathways (**Figure 4A**). Together, these findings indicate that the transcriptomic state space comprises biologically distinct programs and highlight specific modules with greater clinical relevance. Detailed correlation outputs for each module are present in **Supplementary Tables 5 and 6**.

### Transcriptomic state space organisation reveals biologically distinct patient groups in LUAD

Tumours within the module-defined state space were organised along a dominant continuous axis, with the transcriptomic risk score strongly aligned to this variation (Spearman ρ = 0.72), linking the principal direction of transcriptomic variation to clinical outcome. Unsupervised clustering of module activity profiles identified three stable patient groups in the TCGA LUAD cohort, and the module score heatmap showed distinct patterns of module activity across these groups, suggesting that tumours occupy structured regions within the transcriptomic landscape (**Figures 4B and 4C**).

Projection into principal component space showed that the three groups occupied distinguishable but partially overlapping regions along the dominant axis, consistent with their organisation within a continuous transcriptomic landscape (**Figure 4D**). Survival outcomes significantly across the groups (log-rank p = 0.0032), with Group 1 showing the poorest outcomes (median 40.6 months), Group 2 intermediate outcomes (53.3 months), and Group 3 the most favourable prognosis (86 months) (**Figure 4E**).

The groups also displayed distinct genomic and clinical enrichment patterns. Group 1 was enriched for TP53 loss-of-function alterations, higher pathological stage, and current smoking status, whereas Group 2 showed enrichment for activating EGFR mutations and reduced smoking prevalence. Group 3 was associated with reduced TP53 alteration burden and enrichment of STK11 wild-type status. Complete enrichment details are available in **Table 3.Transcriptomic modules reflect tumour–immune–stromal ecosystem organisation**

**Table 3.** Genomic and clinical enrichment patterns associated with transcriptomic patient groups in the TCGA LUAD cohort, with Benjamini-Hochberg-corrected p-values (p-adj).

| Group | Feature | Enrichment (Odds Ratio) | p-adj |
| --- | --- | --- | --- |
| 1 | TP53_LOF mutation | 3.12 | 1.36e-09 |
| 1 | TP53_WT | 0.28 | 8.12e-11 |
| 1 | Activating EGFR mutations | 0.32 | 0.002 |
| 1 | Sex (Male) | 1.58 | 0.03 |
| 1 | Sex (Female) | 0.63 | 0.03 |
| 1 | Pathological stage I | 0.49 | 5.24e-04 |
| 1 | Pathological stage III | 1.88 | 0.03 |
| 1 | Smoking status (Current) | 2.51 | 2.8e-04 |
| 2 | TP53_LOF mutation | 0.4 | 5.29e-06 |
| 2 | TP53_WT | 2.75 | 1.05e-06 |
| 2 | Activating EGFR mutations | 3.24 | 2.1e-04 |
| 2 | Pathological stage II | 0.5 | 0.03 |
| 2 | Smoking status (Current) | 0.35 | 9.3e-04 |
| 3 | TP53_LOF mutation | 0.62 | 0.049 |
| 3 | TP53_WT | 1.64 | 0.049 |
| 3 | STK11_LOF mutation | 0.25 | 0.03 |
| 3 | STK11_WT | 4.0 | 0.03 |

The module-defined transcriptomic structure was evaluated at cellular resolution using a primary LUAD single-cell RNA-seq dataset (GSE131907; n = 15 patients, 57,155 cells) spanning epithelial (tumour), immune, and stromal compartments. Modules 2, 5, and 8 were examined specifically to assess cross-compartment organisation, as they represented the most clinically relevant transcriptomic programmes based on independent prognostic associations (**Figure 4C**). Cell-type– resolved pseudobulk profiles were generated, and module activity scores were projected onto tumour, immune, and stromal compartments using the same GSVA-based framework applied to bulk data (**Figures 5A and 5B**).

**Figure 5.**
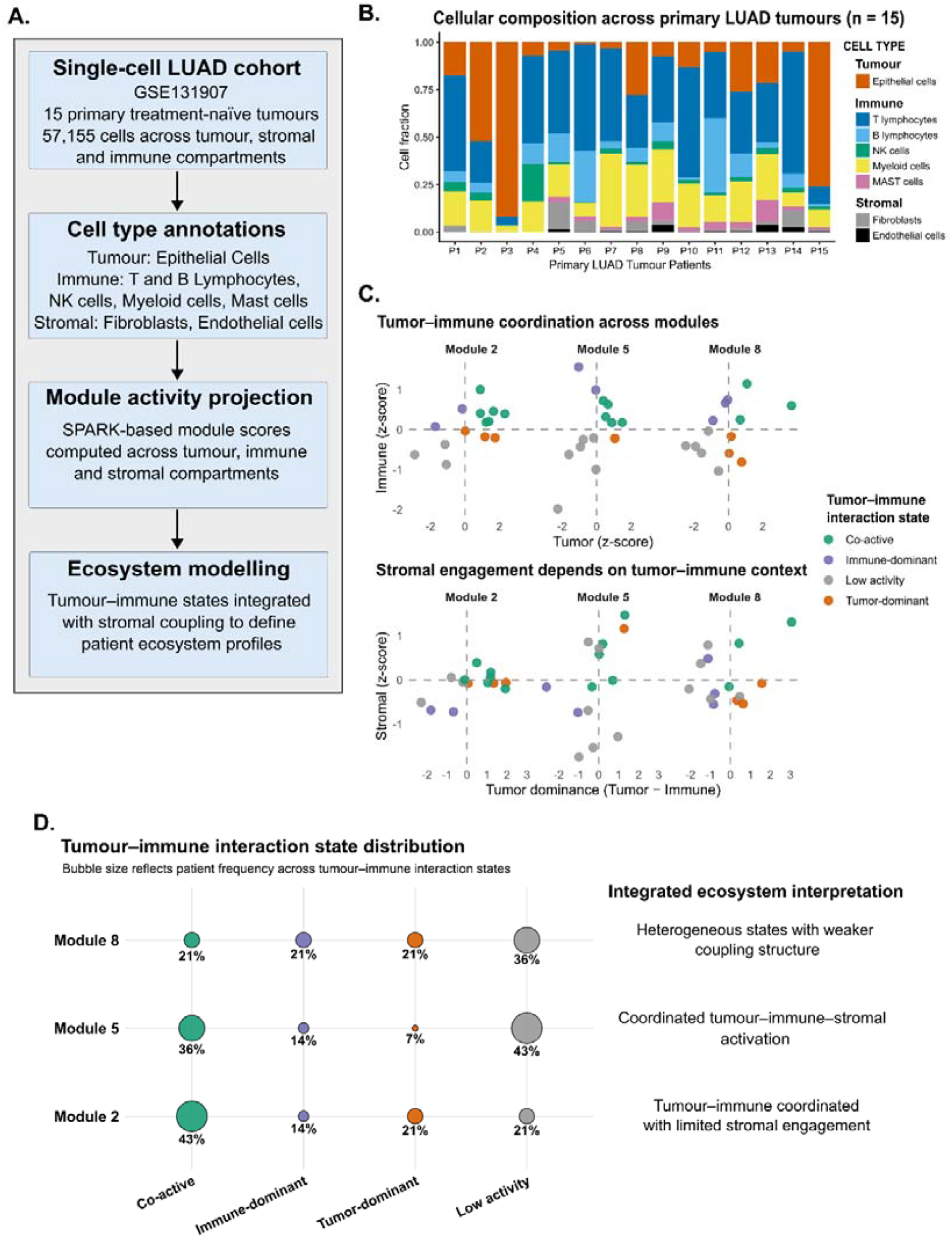
Single-cell evaluation of the tumour–immune–stromal ecosystem. **(A)** Overview of the single-cell LUAD cohort, cell-type annotation strategy, module activity projection workflow, and ecosystem modelling framework. **(B)** Relative tumour, immune, and stromal cell compositions across the 15 primary LUAD tumours in the single-cell cohort. **(C)** Tumour–immune coordination and stromal engagement patterns across Modules 2, 5, and 8 using compartment-level module activity scores. **(D)**Distribution of tumour–immune interaction regimes across Modules 2, 5, and 8, illustrating module-specific ecosystem interaction states and stromal coupling behaviour.

Marked inter-patient variability was observed in the tumour, immune, and stromal cell compositions. Module activity was mapped into the tumour–immune coordinate space using z-score-normalised compartment-level scores. Interaction states were defined by the sign of tumour and immune activity, yielding four regimes: co-active (Tumour ≥ 0, Immune ≥ 0), tumour-dominant (Tumour ≥ 0, Immune < 0), immune-dominant (Tumour < 0, Immune ≥ 0), and low-activity (Tumour < 0, Immune < 0). These regimes were unevenly distributed across modules (**Figure 5C**).

Modules 2, 5, and 8 exhibited distinct interaction patterns across tumour, immune, and stromal compartments. Module 2 was enriched for tumour-dominant and low-activity states, with consistently low stromal activity, indicating limited engagement of non-tumour compartments. Module 5 showed a higher proportion of co-active states and increasing stromal activity with tumour–immune balance, consistent with coordinated activation across compartments. In contrast, Module 8 displayed a heterogeneous distribution across tumour–immune regimes and a weak, inconsistent association with stromal activity, reflecting reduced cross-compartment coordination (**Figure 5D**).

These results indicate that the module-defined transcriptomic structure reflects context-dependent organisation of the tumour–immune–stromal ecosystem. While limited by sample size, this analysis provides orthogonal biological support for the module-defined architecture, linking bulk transcriptomic organisation to tumour– immune–stromal interactions at cellular resolution.

## Discussion

Transcriptomic heterogeneity in LUAD is well established and reflects a complex, high-dimensional landscape of molecular variation across patients, which can complicate biological interpretability and clinical translation [1]. In this study, we showed that this complexity is not arbitrary but is organised into a stable, low-dimensional structure defined by coordinated pathway programs. Using the SPARK framework, multimodal pathway activity patterns were resolved into a small number of modules that capture higher-order transcriptomic organisation. These define a compact representation of each tumour as a transcriptomic state, demonstrating that inter-patient variability can be explained by coordinated shifts across biological programs rather than independent gene-level changes.

The modules identified here suggest that heterogeneity in LUAD is organised along a small number of biologically meaningful dimensions rather than discrete molecular categories [7]. For example, proliferative programmes and tumour suppressor– associated pathways show opposing behaviour, indicating that tumours vary along a spectrum of growth activity. Immune-related modules reflect differences in inflammatory signalling and activation across patients, highlighting variability in tumour–microenvironment interactions. In addition, metabolic, chromatin, and remodelling-associated programs capture other independent aspects of tumour biology.

Integration of module activities into a composite risk score highlighted their direct prognostic relevance in LUAD. The risk score was significantly associated with overall survival. It remained independently predictive after adjustment for clinical variables (HR = 2.66), reflecting that it captures information beyond established clinicopathologic factors such as pathological stage. Survival differences were also observed between high- and low-risk early-stage (I/II) groups, further supporting the risk score’s independent prognostic value. While the transcriptomic model alone showed moderate performance (C-index = 0.635), integration with clinical covariates improved discrimination (C-index = 0.711).

Building on the prognostic relevance of the risk score, the coordinated variation across modules organises tumours along a continuous transcriptomic space. Although clustering identified three patient groups, these occupied partially overlapping regions within the module-defined space and did not represent fully distinct categories. Instead, individual tumours are distributed along a continuum of module activity patterns, reflecting gradual differences in underlying tumour biology across patients. This is supported by the strong correlation between the risk score and PC1 (Spearman ρ = 0.72), suggesting that the risk score highlights a major axis of transcriptomic variation. These findings indicate that tumour heterogeneity may be better represented as a continuous organisation of transcriptomic states, where clinical risk reflects position along this spectrum rather than membership in discrete subtypes, consistent with prior observations that molecular organisation can transcend rigid categorical classifications [26].

Module activity projected onto tumour, immune, and stromal compartments using single-cell data demonstrates that the programs reflect coordinated cross-compartment organisation rather than tumour-intrinsic signals alone. The contrasting behaviour of Modules 2, 5, and 8 indicates that tumours occupy states defined by the extent of tumour–immune–stromal coordination, spanning tumour-dominant to integrated ecosystem-level activity. This provides direct evidence that the transcriptomic state framework successfully represents a system-level organisation of the tumour microenvironment. This suggests that variation in bulk transcriptomic profiles reflects underlying differences in cellular interactions, linking the module-based state space to biologically meaningful ecosystem organisation [2].

While these results establish the transcriptomic state as a systems-level representation of tumour organisation, several limitations remain. The analyses are based on retrospective cohorts and require prospective validation for clinical translation. Module definitions from bulk RNA-seq capture composite signals across cell types, and the single-cell analysis is limited in sample size. Nevertheless, this framework enables an investigation of how coordinated pathway programs and tumour–microenvironment interactions might relate to therapeutic response and resistance. Integration with spatial and multi-omic data may further refine tumour state characterisation and improve clinical relevance by linking states to therapy response and resistance.

## Data and Code Availability

The complete analysis workflow, including source code, processed data objects, and scripts required to reproduce the results presented in this study, is available at https://github.com/CGnTLab/SPARK.

Bulk RNA-sequencing and clinical data for the TCGA-LUAD and CPTAC-LUAD cohorts were obtained from publicly accessible repositories through cBioPortal and the Genomic Data Commons (GDC). Single-cell RNA-sequencing data were obtained from the Gene Expression Omnibus (GEO) under accession GSE131907.

## Author Contributions

Rishabh Kulkarni (Data curation [lead], Formal analysis [lead], Methodology [lead], Visualisation [lead], Software [equal], Writing-original draft [lead], Writing-review & editing [equal]), Avik Sengupta (Software [equal]) and Rahul Kumar (Project administration [lead], Supervision [lead], Writing-review & editing [equal]).

## Acknowledgements

We acknowledge the Indian Institute of Technology Hyderabad for providing research infrastructure. We thank Dr Pratik Chandrani for his critical scientific inputs.

---

**Supplementary Figure 1. Additional evaluation of transcriptomic risk model robustness. (A)** Elastic-Net feature stability across 200 bootstrap iterations. The dashed line indicates the predefined feature selection threshold. **(B)** Permutation importance analysis showing reduction in model C-index following module-wise feature shuffling. **(C)** Kaplan–Meier survival analysis for high-risk and low-risk groups within early-stage (stage I/II) LUAD patients (log-rank p = 0.00067).

