## Supplementary Figure 1 for "SPARK: A Systems-level Computational Framework for Reconstructing Transcriptomic State Organisation in Lung Adenocarcinoma"

**Supplementary Figure 1. Additional evaluation of transcriptomic risk model robustness. (A)** Elastic-Net feature stability across 200 bootstrap iterations. The dashed line indicates the predefined feature selection threshold. **(B)** Permutation importance analysis showing reduction in model C-index following module-wise feature shuffling. **(C)** Kaplan–Meier survival analysis for high-risk and low-risk groups within early-stage (stage I/II) LUAD patients (log-rank p = 0.00067).

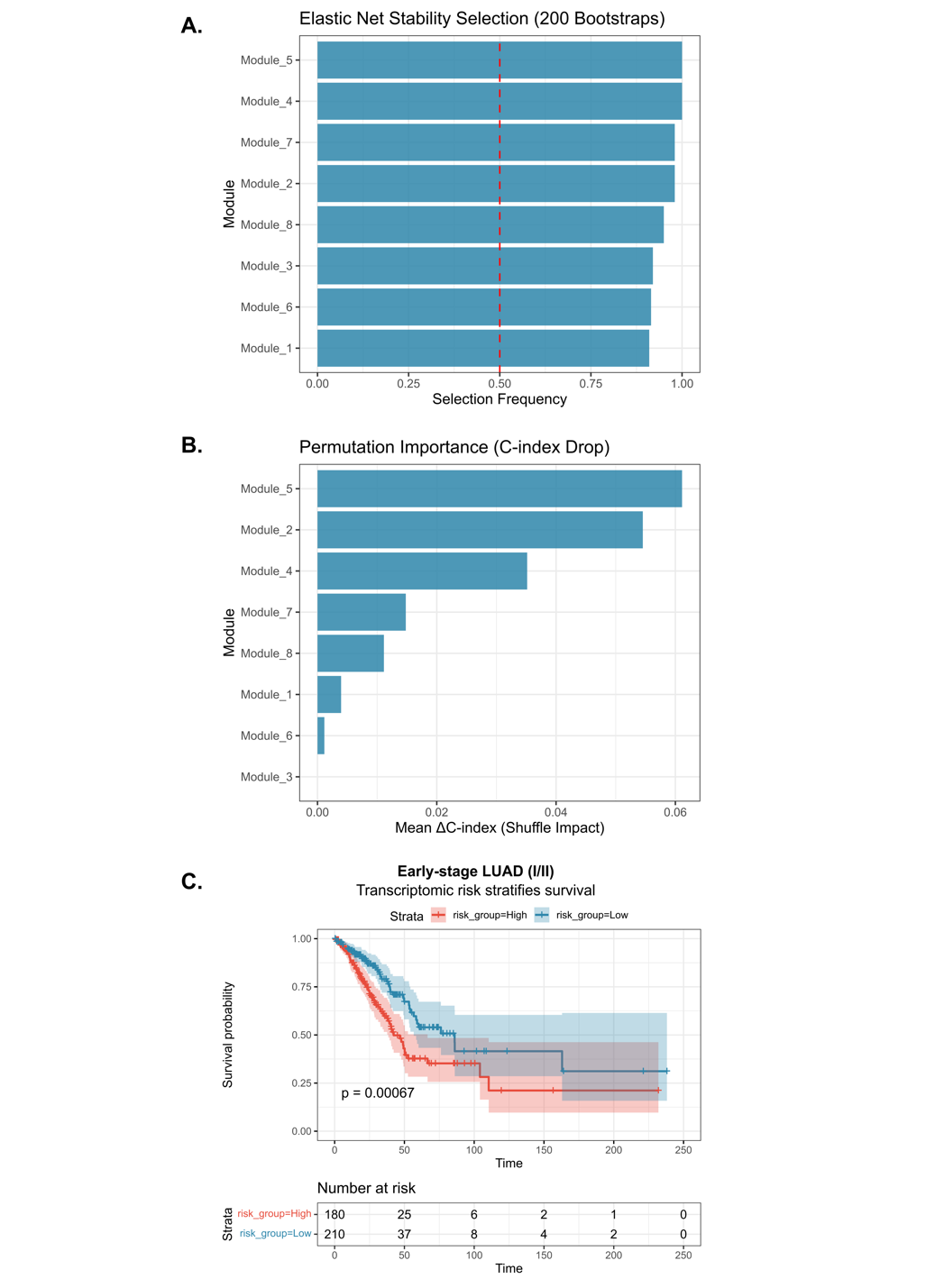
